# GDSCTools for Mining Pharmacogenomic Interactions in Cancer

**DOI:** 10.1101/166223

**Authors:** Thomas Cokelaer, Elisabeth Chen, Francesco Iorio, Michael P. Menden, Howard Lightfoot, Julio Saez-Rodriguez, Mathew J. Garnett

## Abstract

**Motivation:** Large pharmacogenomic screenings integrate heterogeneous cancer genomic data sets as well as anti-cancer drug responses on thousand human cancer cell lines. Mining this data to identify new therapies for cancer sub-populations would benefit from common data structures, modular computational biology tools and user-friendly interfaces.

**Results:** We have developed GDSCTools: a software aimed at the identification of clinically relevant genomic markers of drug response. The Genomics of Drug Sensitivity in Cancer (GDSC) database (www.cancerRxgene.org) integrates heterogeneous cancer genomic data sets as well as anti-cancer drug responses on a thousand cancer cell lines. Including statistical tools (ANOVA) and predictive methods (Elastic Net), as well as common data structures, GDSCTools allows users to reproduce published results from GDSC, to analyse their own drug responses or genomic datasets, and to implement new analytical methods.

## 1 Introduction

Cancers occur due to genetic alterations in cells accumulated through the lifespan of an individual. Cancers are genetically heterogeneous and as a consequence patients with similar diagnoses may vary in their response to the same therapy. The path towards precision cancer medicine requires the identification of specific biomarkers, such as genetic alterations, allowing effective patient selection strategies for therapy.

Large-scale pharmacological screens such as the Genomics of Drug Sensitivity in Cancer (GDSC) (Garnett *et al.*, 2012) and Cancer Cell Line Encyclopaedia (CCLE) projects (Barretina *et al.*,2012) have been used to identify potential new treatments and to explore biomarkers of drug sensitivity in cancer cells. In particular, the GDSC project releases database resources periodically (www.cancerRxgene.org) (Yang *et al.*,2013). A recent installment of this resource (version 17, v17 hereafter) includes cancer-driven alterations identified in 11,289 tumors from 29 tissues across 1,001 molecularly annotated human cancer cell lines, and cell line sensitivity data for 265 anti-cancer compounds. A systematic identification of clinically-relevant markers of drug response uncovered numerous alterations that sensitize to anti-cancer drugs (Iorio *et al.*,2016). Here, we present GDSCTools, a Python library that allows users to perform pharmacogenomic analyses as those presented in (Iorio *et al.*,2016). Our software complements an existing tool (Smirnov *et al.*,2016) by giving access to the full GDSC dataset and providing a powerful platform for statistical analyses and data mining through visualization tools. In the following, we briefly describe the GDSCTools features, including common data structures, statistical tools and machine learning approaches.

## 2 Data formats and data wrangling tools

The GDSC database provides large-scale genomics and drug sensitivity datasets. The drug sensitivity dataset contains dose-response curves (e.g., cell viability for 5 - 9 drug concentrations) which can be used to derive drug sensitivity indicators (Vis *et al.*,2016; Garnett *et al.*,2012), such as the half-maximal inhibitory concentration (IC_50_) or the area under the curve (AUC) (See Fig. 1-A). In GDSCTools, IC_50_ indicators are encoded as a *Nc × N_d_* matrix, where *Nc* is the number of cell lines labeled with their COSMIC identifier (http://cancer.sanger.ac.uk/cosmic) and *N_d_* is the number of drugs. For a given drug, we denote with **Y_d_** the vector of IC_50_s across the *Nc* cell lines. The genomic feature dataset **X** is also encoded as a *Nc ×N_f_* matrix, where *N_f_* is the number of genomic features. In addition to a subset of the v17 data files available in GDSCTools, users can also retrieve additional data sets online (e.g., methylation data, copy number variants, etc.). Database-like queries can be used to extract and use specific features (e.g., only gene amplifications or deletions). These database-like functionalities are part of the *OmniBEM* builder (see supplementary section).

**Fig. 1.**
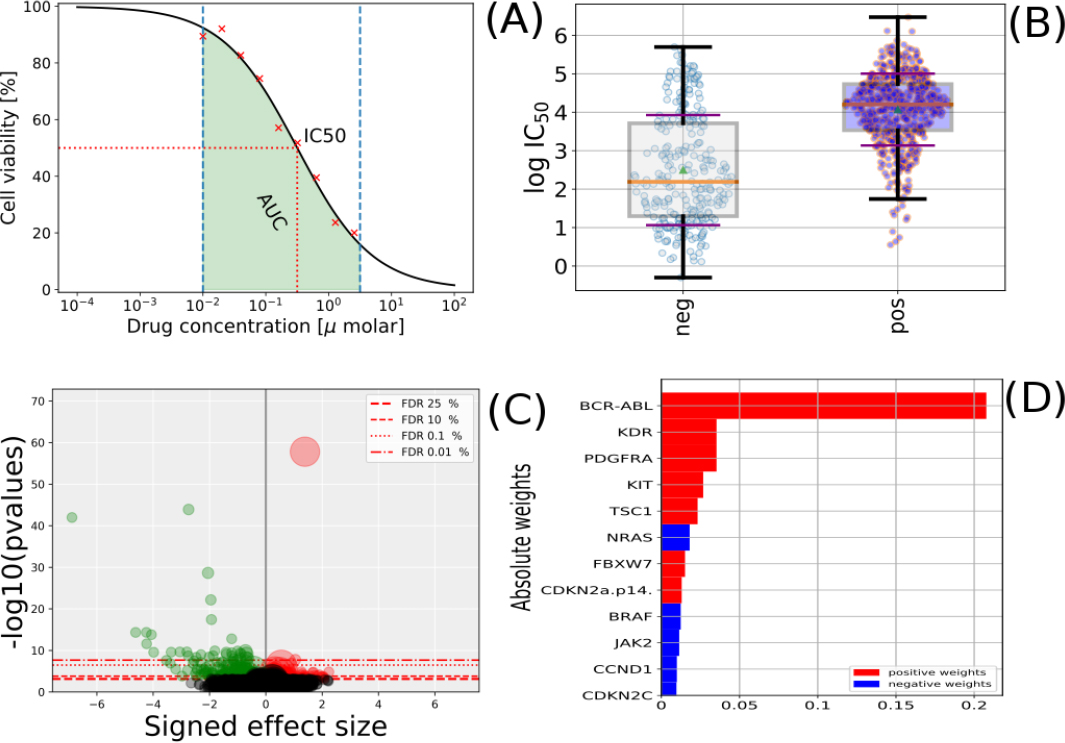
Panel A: drug response (cell viability versus drug concentrations) and derived drug response metrics (AUC and IC_50_ s). Panel B: distribution of IC_50_ s in response to a given drug across a dichotomy of cell lines induced by the status of a genomic feature. Panel C: p-values from an ANOVA analysis versus signed effect sizes (all drug-genomic feature interactions). Panel D: weights distributions resulting from training a sparse linear regression model of a given drug response using all the genomic features.

## 3 Data analysis tools

Using GDSCTools, genomic features can be investigated as possible predictors of differential drug sensitivity across screened cell lines. The statistical interaction **Y_d_** *∼* **X** between drug response and genomic features can be tested within a sample population of cell lines from the same cancer type with a t-test. However, to account for possible confounding factors (including the tissue of origin, when performing pan-cancer analyses) a more versatile analysis of variance (ANOVA) is implemented. In this model the variability observed in **Y_d_** is first explained using the tissue covariate, subsequently using additional factors (e.g., microsatellite instability denoted by MSI), and finally by each of the genomic features in *X* (one model per feature). This can be mathematically expressed as *Y_d_ ∼ C*(*tissue*) + *C*(*M SI*) + *…* + *f eature*, where the *C*() operator indicates a categorical variable. An ANOVA test is performed for each combination of drug and genomic feature (Fig. 1-B). Outcomes of this large number of tests (*N_d_ × N_f_*) are corrected for multiple hypothesis testing using Bonferroni or Benjamini-Hochberg corrections. To account for p-value inflations due to differences in sample sizes, the effect sizes of the tested statistical interactions (computed with the Cohen and Glass models) are also included (Fig. 1-C).

Unlike the ANOVA analysis that is performed on a one drug / one feature basis, linear regression models assume that drug response can be expressed as a linear combination of the status of a set of genomic features. GDSCTools includes 3 linear regression methods: (i) Ridge, based on an L2 penalty term, which limits the size of the coefficient vector; (ii) Lasso, based on an L1 penalty term, which imposes sparsity among the coefficients (i.e., makes the fitted model more interpretable); and (iii) Elastic Net, a compromise between Ridge and Lasso techniques with a mix penalty between L1 and L2 norms (see supplementary for details). The linear regression methods require optimisation of an *α* parameter (mix ratio between L1 and L2). This is performed via a cross validation to avoid over-fitting. The best model is determined using as objective function the Pearson correlation between predicted and actual drug responses on the training set. The final regressor weights are outputted as shown in Fig. 1-D. Significance of the final selected models is computed against a Monte Carlo simulated null model.

## 4 Implementation and future directions

GDSCTools is available on http://github.com/CancerRxGene/gdsctools. It is fully documented on http://gdsctools.readthedocs.io. Pre-compiled versions of the library are available on https://bioconda.github.io/. GDSCTools can be used via standalone applications to analyse a user defined set of drugs (and genomic features) and assemble the results in an HTML report. We also provide solutions based on the Snakemake framework (Köster and Rahman, 2012) to parallelize the analysis on distributed cluster farm architectures such as LSF or SLURM (see supplementary data). Besides analysis of pharmacogenomic datasets, GDSCTools can provide the framework for discovering new biomarkers through integration/mining of novel and heterogeneous data sets, including pharmacological, RNA interference or increasingly available genetic screens (e.g., CRISPR), alternative drug response metrics (e.g., AUC), or implementing new analytical tools. The augmentation of genomic features with information obtained from online web services (Cokelaer *et al.*, 2012) like pathway enrichment (e.g., via OmniPath (Turei *et al.*, 2017)) will further extend functionality and usefulness of GDSCTools.

## 5 Supplementary

### 5.1 Code and Installation

GDSCTools source code is available on GitHub website at http://www.github.com/cancerRxgene/gdsctools and a pre-compiled version is available on Bioconda channel. It has an extended documentation hosted on http://gdsctools.readthedocs.io. We have also included a large set of functional tests to assess results’ reproducibility. Changes made to the code are tested automatically via the Travis continuous integration framework so that changes that affect the analysis or normal behaviour may be caught early. Finally, GDSCTools makes use of existing open source libraries such as scikit-learn (Pedregosa *et al.*,2011) for the machine learning tools, and statsmodels (Seabold *et al.*,2010) for advanced statistical analysis.

GDSCTools can be installed from the source code. However, we recommend using the Anaconda framework. Information can be found on https://www.continuum.io/downloads. Once the software is installed, an executable called *conda* provides pre-compiled versions of many scientific libraries. In addition, GDSCTools itself is exposed on one of the Anaconda channel called **Bioconda**. Consequently, having Anaconda pre-installed makes the installation of GDSCTools easier and quicker. The commands needed to select the Anaconda channel to be used (once for all) are the following ones:

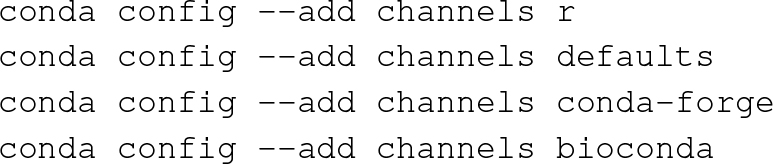

Then, GDSCTools can be installed as follows:

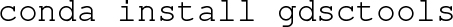

This will take care of all dependencies required by GDSCTools. Further details can be found in the http://gdsctools.readthedocs.io website (installation section).

### 5.2 Data

#### 5.2.1 IC_50_ indicators

The first data object that GDSCTools uses by default contains IC_50_ indicators, summarising the effect of drug treatment across a large collection of cell lines using experimental protocols detailed in (Garnett *et al.*,2012; Iorio *et al.*,2016). These indicators were derived by applying a curve-fitting algorithm to raw cell counts data, via a multilevel mixed model (Vis *et al.*,2016). These (or any other user defined drug response indicators) must be stored in a CSV file, which can be optionally compressed (gzip format). In this file, the header must contain an entry named COSMIC_ID: this column will contain the COSMIC identifiers of the cell lines, one per line. The following entries should contain drug identifiers (one integer per column). The order of the columns is not relevant. Each row should contain IC_50_s for a given cell line (identified through its COSMIC_ID), across all the tested drugs. Here is an example of the data format for 2 cell lines and 3 drugs

|  |  |  |  |
| --- | --- | --- | --- |
| COSMIC_ID, | 1, | 20, | 40 |
| 111111, | 0.5, | 0.8, | 10 |
| 222222, | 1, | 2, | 10 |

Further details can be found within the GDSCTools on-line documentation http://gdsctools.readthedocs.io/en/master/data.html. Worthy of note is that in this data object, IC_50_s can be replaced by any kind of scalar data (e.g., AUCs). To read the IC_50_s file shown above, the following commands should be used (assuming that the object is saved into a file named *ic50.csv*):

**Fig. 2.**
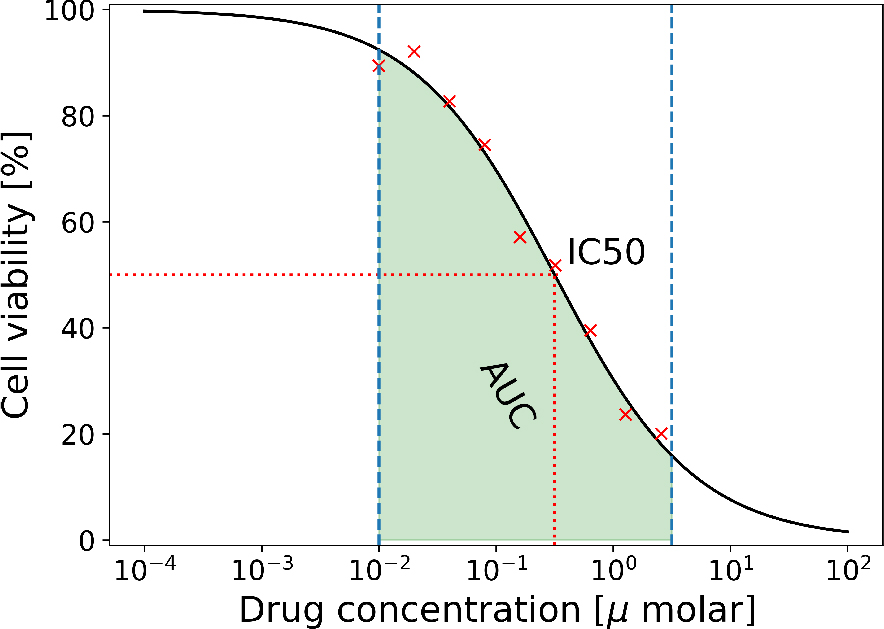
Drug response (cell viability upon exposure to different drug concentrations) and derived drug response metrics (AUC and IC_50_).

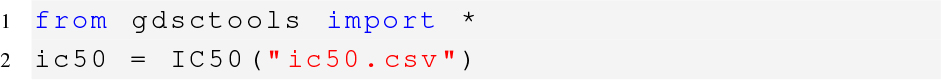

This allows the data to be accessed as a DataFrame and used with various descriptive statistics and plotting functions.

#### 5.2.2 Genomic features

The second data set required by GDSCTools is the *Genomic Features* data set. All the implemented analyses are performed on the cell lines included in both the IC_50_s and the *Genomic Features* data object and this intersection is determined at the level of COSMIC identifiers. As a consequence, cell lines that are not included in both matrices will be discarded. In addition to the COSMIC identifiers, the *Genomic Features* file should contain the following columns: TISSUE_FACTOR, MSI_FACTOR and MEDIA_FACTOR that can be used in the ANOVA or linear regression models as explained hereafter. All remaining columns should refer to individual genomic features, whose status (positive or negative) in a generic *i−*th cell line should be indicated with a binary entry in the *i−*th line.

An example is reported below.

|  |  |  |  |
| --- | --- | --- | --- |
| COSMIC_ID, | TISSUE_FACTOR, | MSI_FACTOR, | BRAF_mut |
| 111111, | lung_NSCLC, | 1, | 1 |
| 222222, | prostate, | 1, | 0 |

To read this *Genomic Features* object saved for example in the *gf.csv* file, the following commands should be executed:

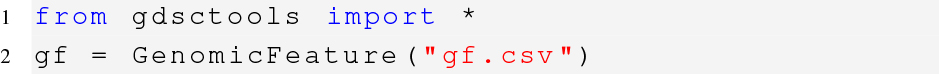

The genomic feature data is accessible as a DataFrame with plotting and statistical capabilities.

#### 5.2.3 Data retrieval and data wrangling

In GDSCTools, we embedded several data sets either for testing purposes or to serve as full scale examples. One such type of data is related to the version 17 (v17) of the GDSC database used in (Iorio *et al.*,2016). A subset of the genomic features and IC_50_s used in (Iorio *et al.*,2016) are provided inside GDSCTools, which includes IC_50_s of 265 drugs across 988 cell lines. In parallel, a *Genomic Features* file encompassing the status of 677 genomic features (copy number alteration and cancer gene mutations) on the same set of cell lines is provided. Alternatively, GDSCTools contains built-in functions to retrieve and to analyse more data from the GDSC database. This currently encompasses data sets including 29,214 gene variants, 2,436 copy number variations (deletion and amplification) and 10,581 differentially methylated gene promoters across 1,002 cancer cell lines. For instance, the following code downloads all GDSC data from (Iorio *et al.*,2016) locally in a data directory (line 3). One can filter the data to keep only a sub set of *Core Genes* used in the published analysis.

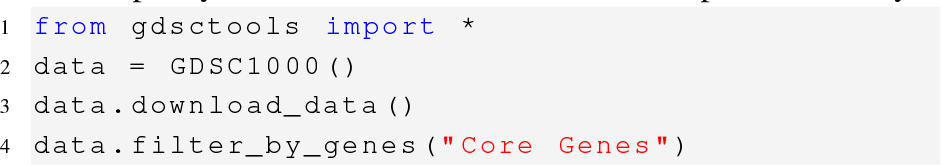

More generally, we provide the *OmniBEM* Builder module that allows the user to merge different levels of annotations from the GDSC web-portal into a single gene-level view that merges together different types of alterations (for example mutations and copy number amplifications involving the same gene). Users can additionally specify which sets of genomic annotations to integrate as well as upload and integrate their own sets of genomic annotations.

### 5.3 ANOVA

#### 5.3.1 Details

GDSCTools implements functions to perform a systematic analysis of variance (ANOVA) to identify statistically significant interactions between genomic features and drug responses. To this aim IC_50_s and *Genomic Features* data object must be created first (as explained in the previous section).

The implemented model is fully detailed in (Iorio *et al.*,2016) and (Garnett *et al.*,2012). Briefly, for each drug a drug response vector is assembled consisting of *n* IC_50_s values, derived from treating *n* cell lines with the drug under consideration, as explained in the previous sections. The implemented model is linear with no interaction terms, dependent variables represented by the described vector and independent factors including tissue type, and screening medium (for the pan-cancer analysis only), microsatellite instability status (for the cancer types with positive samples for this feature) and the status of a genomic feature. For all the tested gene-drug associations, an indication of their effect size is estimated considering the pooled standard deviation of the analysed IC_50_s population (Cohen’s d), or the individual standard deviations (quantified through two different Glass deltas), for the IC_50_s populations of the cell lines that are respectively positive or negative for a given genomic feature. P-values and all other statistical scores are obtained from the fitted models. A genomic-feature/drug pair is tested only if at least *n* cell lines are contained in the two sets resulting from the dichotomy induced by the status of the considered genomic-feature (for example at least 3 positive cell lines and at least 3 negative cell lines), and *n* can be defined by the user.

The resulting p-values are corrected (all together those obtained in the pan-cancer analysis and on a cancer type basis those obtained in a given cancer-specific analysis), with a user-chosen method among Bonferroni (Bonferroni,1935) or Benjamini-Hochberg (Benjamini-Hochberg *et al.*,1995).

#### 5.3.2 Examples

From GDSCTools Python library, which is fully documented on http://gdsctools.readthedocs.io, users can read the IC_50_s and *Genomic Features* files, perform the analysis and create HTML reports highlighting the identified significant interactions and meaningful models. Here is the code to perform these tasks:

**Fig. 3.**
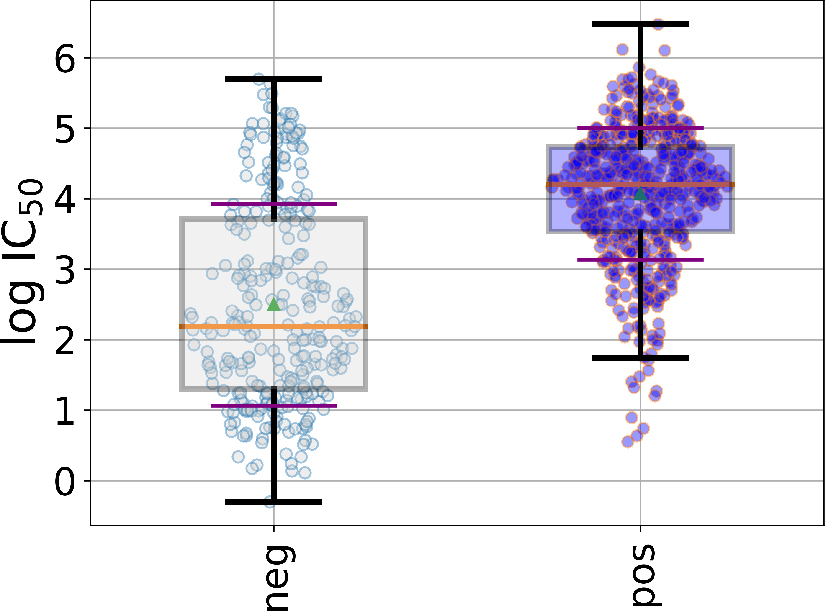
Distribution of IC_50_ s for a given drug and genomic feature. The wild type is shown on the left and the IC_50_ s corresponding to the mutated cell lines are shown on the right.

**Fig. 4.**
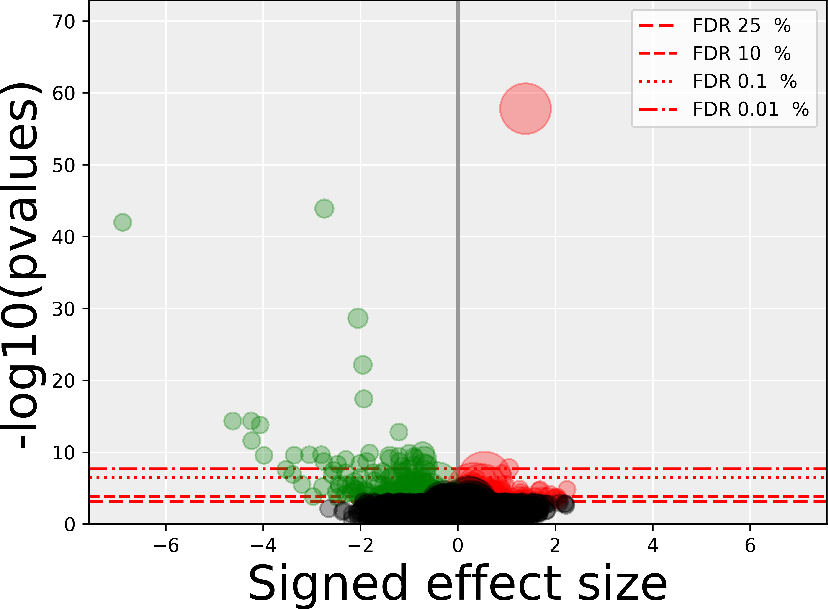
Summary of the ANOVA analysis across all drugs and all features (ADAF). p-values from the ANOVA analysis are shown versus signed effect sizes. Red horizontal lines indicates several false discovery rate (FDR) that is the rate of type I errors in null hypothesis testing when conducting multiple testing corrections (such as Bonferroni correction).

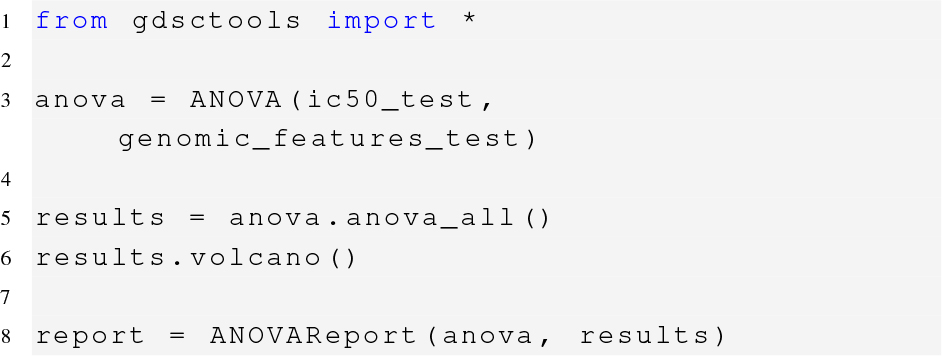

On line 1, the entire library is loaded. On line 3, the ANOVA class is called. The first and second arguments are the IC_50_s and *Genomic Features* files. Here, we use two test files embedded in GDSCTools (only 11 drugs, 47 mutations, 10 tissues). This is for test purposes and can be replaced with other files. On line 5, we run the ANOVA analysis. This may be time consuming depending on the number of drugs and genomic features, although this example would take only a few seconds on a typical desktop computer. Once the analysis is completed, users can look at results with multiple visualisation routines. For example in the form of volcano plots 4 that shows the p-values versus signed effect size for each combination of drug and genomic features. An alternative to the Python library is to use a standalone application from a shell. It is named *gdsctools_anova* and has its own online help. Consider this code:

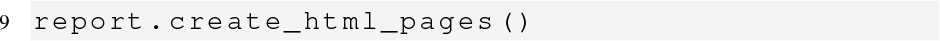

This performs the ANOVA analysis on each drug and genomic feature of the version 17 of the GDSC data. Then, it creates HTML reports that can be browsed to inspect the significant associations more closely. The two data files can be found within the GDSCTools library or downloaded as follows

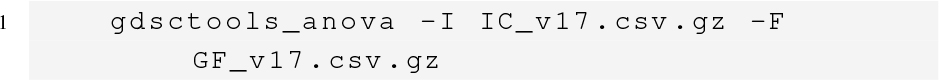

### 5.4 Regression analysis

The Elastic Net model is a linear regression model trained with L1 and L2 priors as regularizer. The objective function to minimize is defined as

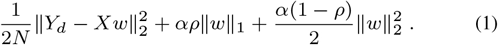

Where *Y_d_* as defined before contains the IC_50_s for all cell lines for given drug and *X* contains the genomic features for the same cell lines. Here we will use the notations used in the scikit-learn library (Pedregosa *et al.*,2011). The mixing parameter *ρ* (with 0 *≤ ρ ≤* 1) controls the combination of L1 and L2 penalties. For *ρ* = 1 the penalty is an L1 penalty (Lasso) while for *ρ* = 0 we have an L2 penalty (Ridge). In the Elastic Net analysis, we fix *ρ* = 0.5 but it can be changed by the user. The equation above allows for learning a sparse model where few of the weights are non-zero like Lasso, while still maintaining the regularisation properties of Ridge. Elastic-net is useful when there are multiple features which are correlated with one another. Lasso is likely to pick one of these at random, while elastic-net is likely to pick both.

Before proceeding with an analysis, we need to minimize the function and optimize the alpha parameter. In order to avoid over-fitting, we hold out part of the available data as a test set and perform a cross validation on a training set. When performing a *k* folds, we train the model with k-1 of the training data and the resulting model is validated on the remaining part of the data. The performance measure reported by *k*-fold cross validation is the average of the values computed on the *k −* 1 models. The metric used to select the best model is the Pearson correlation between predicted and actual drug responses. We scan the range of *α* parameter and select the best *α*. In Fig. 6 we show the Pearson coefficient along the log of an *α* parameter.

**Fig. 6.**
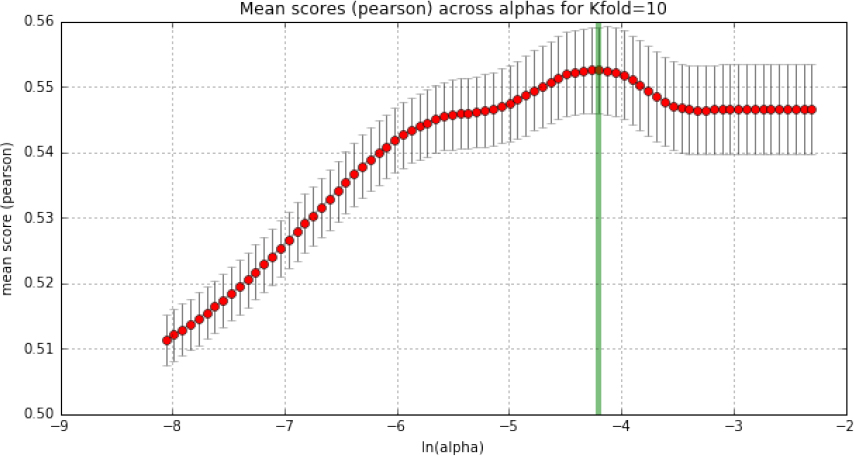
Tuning of the *α* parameter of a linear regression model. Using a 10-folds cross validation, for a given drug and a set of genomic features, we scan the *α* parameter space to obtain the best *α* that maximises the Pearson correlation (indicated by the green vertical line).

### 5.5 Running analysis with Snakemake pipelines

In parallel computing, an embarrassingly parallel problem is one where little or no effort is needed to separate the problem into a number of parallel tasks. In GDSCTools, each drug can be analyzed independently of the others. The analysis is therefore an embarrassingly problem. In GDSCTools, developers can write their own pipelines and run analysis locally, however, we also provide Snakemake pipelines that can be easily run on various clusters (e.g., LSF, SLURM).

#### 5.5.1 Linear models

The initialisation of the pipeline works as follows:

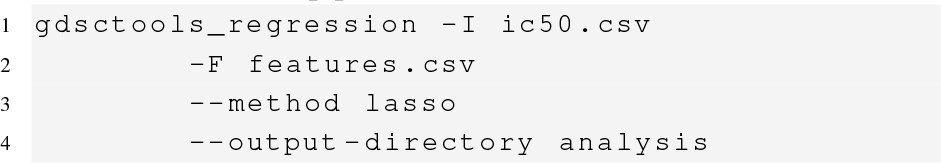

This command creates a directory called *analysis* where a pipeline encoded with the Snakemake framework (Köster and Rahman,2012) is copied. The pipeline filename is *regression.rules*. In addition, a configuration file named *config.yaml* is also provided. A snapshot of the pipeline workflow is shown in Fig.5. This is a simple example with 4 drug responses. Of course, real case examples would include hundreds of them.

**Fig. 5.**
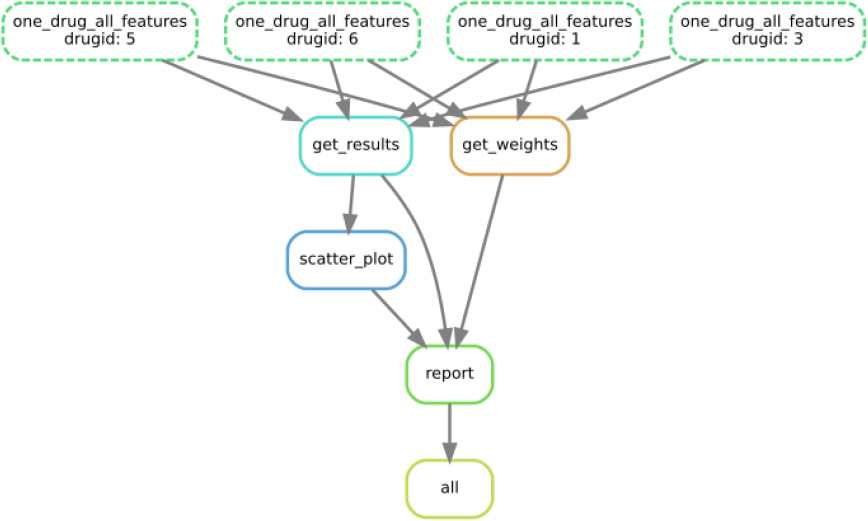
Directed acyclic graph representation of the sparse linear regression pipeline (e.g., Lasso). Each drug can be analyzed independently of the others. We provide Snakemake pipelines that can be easily run on various clusters (e.g., LSF, SLURM). In this example, the input data sets contains 4 drugs (top layer) that can be analysed at the same time. Once the analysis is over, the get_results and get_weights gather the results. Finally, plots and reports are created. The outcome is a HTML file summarising the analysis.

Each drug is analysed in the same way with a linear model analysis (e.g. Lasso). The results of the analysis as well as images representing the weights are stored in sub directories. Finally, HTML reports are created for each drug and a summary page is also created.

The configuration file is the entry point for the end user who can change some parameters such as the regression method, the number of cross validations to perform or the number of null models to compute to compare the best model obtained with a null hypothesis (where *Y* variable is randomized).

Here is the configuration, which can easily be edited and adapted to your needs.

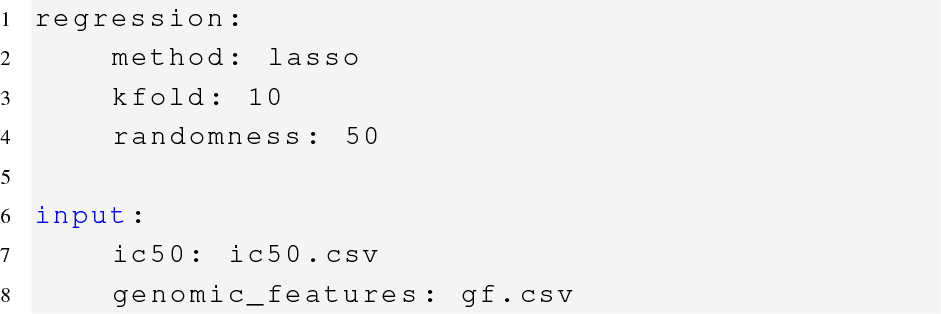

Once the configuration file is available, one can start the analysis as follows. On a local computer (using 4 CPUs):

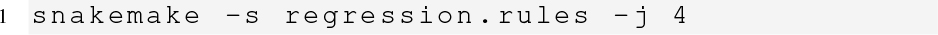

Or on a cluster, you may add the following information (for instance on a SLURM system):

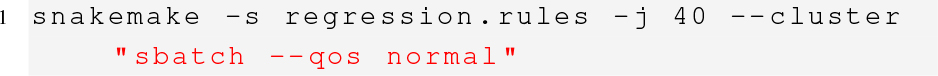

where **-j 40** indicates that we wish to use 40 cores.

